# Model-free Identification of Phenotype-Relevant Variables From Dose Response Data

**DOI:** 10.1101/2023.06.21.545943

**Authors:** Alan Veliz-Cuba, David Murrugarra, Randal Voss

## Abstract

Complex phenotypic changes occur during development and in response to injury and disease. Identifying key regulators of phenotypic change is a shared aim of many different fields of research, including life history, tissue regeneration, and cancer. These examples of phenotypic change involve coordinated changes in cellular behaviors and associated changes in gene expression which if disrupted, can alter and even block completion of the phenotypic transition. Experimental treatments that effectively block the completion of a phenotypic transition can be quantitatively manipulated to identify key regulatory genes through changes in transcriptional dose response. In this paper, we present a “model free” approach to identify “bifurcation-like” behaviors of key regulatory genes by tracking spikes in their transcriptional sensitivities. Genes that exhibit such behavior are predicted to comprise nodes in subnetworks or modules that regulate the phenotypic transition. We applied our method to an in-silico data set where we also studied the impact of noise in the predictions. We also applied the method to a gene expression data that were collected during tail regeneration in axolotls. Our code for gene identification, which can be extended more generally to other component variables of complex phenotypic change, is freely available via the following GitHub site github.com/alanavc/id-vars-from-resp-data.

## 1 Introduction

In multiple areas of biological research it is of interest to identify from a large set of genes which are the most important genes that are associated with a change in phenotype. Dramatic changes in phenotype continue to be a focus of many gene expression studies, including metamorphic transitions that occur during biphasic life histories, injury responses that engender tissue repair and regeneration, and abnormal patterns of growth that lead to organ degeneration and cancer [10, 11, 7, 6]. In many cases, perturbation experiments have been performed to alter or block a phenotypic transition in attempt to decompose regulatory architecture and identify component variables. Pharmaceuticals can be readily delivered to cells, tissues, and organisms for the purpose of altering developmental, cellular, and molecular mechanisms in a dose-dependent manner [12]. Indeed, many data sets have been generated to test the dose dependency of pharmaceuticals on gene expression and phenotypic outcome [1]. Analytical approaches to identify key dose response genes often assume the nature of gene-gene associations within regulatory networks from pre-existing biological data. However, protein interaction and gene co-expression data have not been generated for many organisms; and even for well-studied model organisms, biological inferences from in vitro studies may not translate to in vivo studies [5]. In cases were gene associations and gene regulatory connectivity is unknown, there is need for model-free approaches to identify key genes from dose response experiments.

Here, we explore a model-free approach to identify relevant variables from dose response data. We explore a toy network model to highlight the features and possible limitations of the method. We then validate and measure the accuracy of the approach with an in-silico network model of 24 nodes where we know which variables are directly affected by an external input. Finally, we apply the approach to a gene expression dataset that was collected during axolotl (*Ambystoma mexicanum*) tail regeneration experiments under chemical perturbation.

## 2 Approach

### 2.1 Main idea

Consider a network with unknown connectivity (Fig. 1a) such that some of its variables are responsible for a certain phenotype. The network here can represent a set of genes and the phenotype can be any observable feature. In Fig. 1a we also have an illustration of what that phenotype can be. Under normal conditions, an organism is able to regenerate an appendage when it is cut. We refer to this as *phenotype 1*. In this description, we are interested in the set of variables responsible for this regeneration process. We are interested in using data generated by perturbing the network by an external input, *E*, like a chemical, such that the perturbation causes a change in phenotype when a critical concentration, *E*_0_ is crossed. In our schematic, the organism cannot regenerate an appendage anymore. We refer to this as *phenotype 2*.

**Figure 1:**
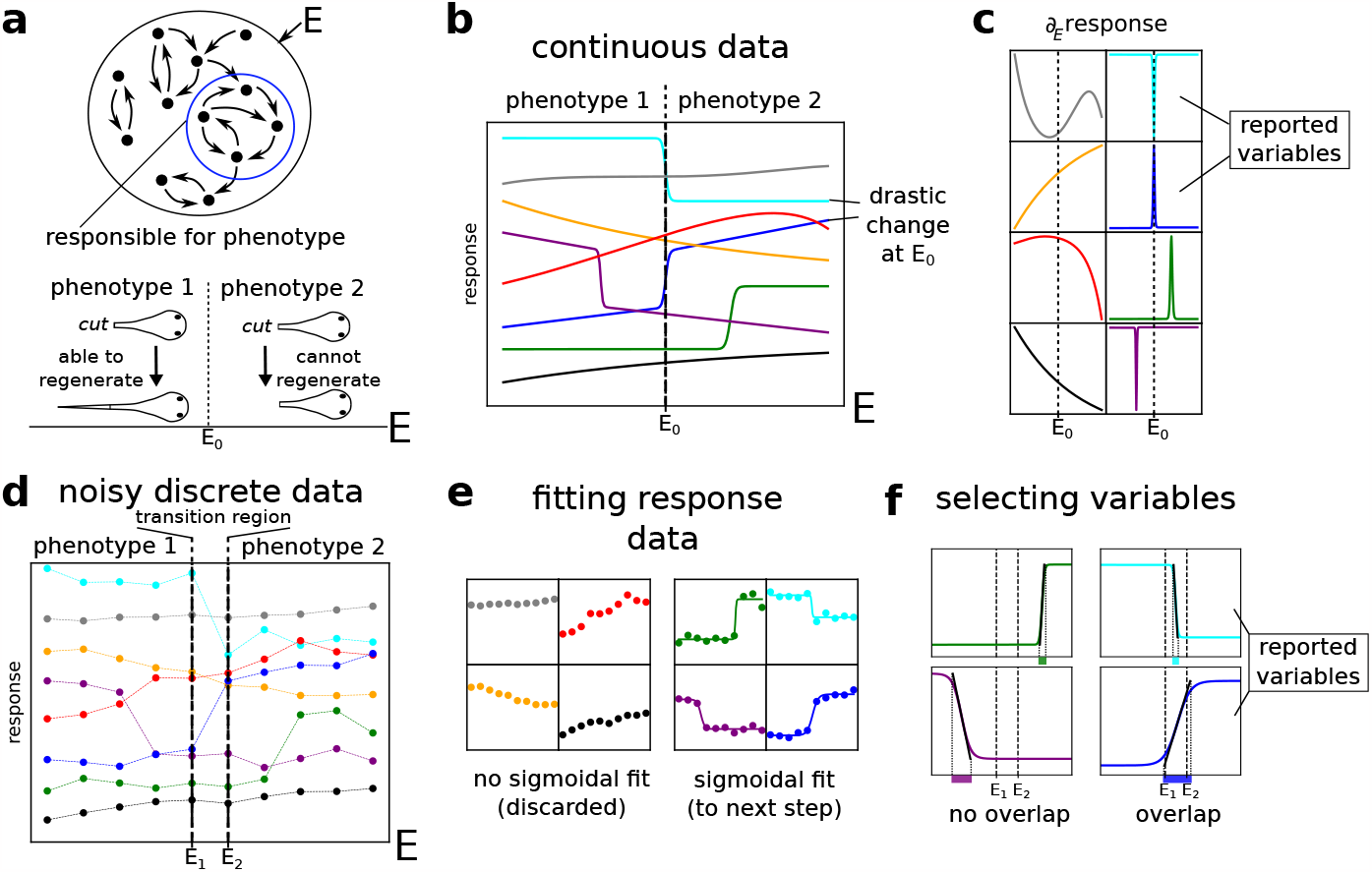
Schematic of the approach. (**a**) Network with subnetwork to be identified. Network is subjected to an external perturbation and exhibits a change in phenotype at *E* = *E*_0_. (**b-c**) **Approach using continuous Data**: (**b**) Different response values are observed for different values of *E*. Potential candidates for the network of interest will have an abrupt change at *E* = *E*_0_, since that is where the phenotypic change is observed. (**c**) The abrupt change is seen as a jump in the sensitivities, 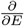 response, at *E* = *E*_0_ (cyan and blue curves). (**d-f**) **Approach using noisy discrete Data**: (**d**) Real data is discrete and subject to noise. The change in phenotype is not observed at a single point, but there are values *E*_1_ and *E*_2_ such that phenotype 1 is observed for *E* ≤ *E*_1_ and phenotype 2 for *E* ≥ *E*_2_. We refer to the interval [*E*_1_, *E*_2_] as the *transition region*. (**e**) Variables that can be fit with a sigmoidal curve are selected for the next step and other variables are discarded. (**f**) We compare the regions where the sigmoidal fits have a steep increase (same color as the curves) with the transition region (interval [*E*_1_, *E*_2_]). If the regions overlap, such variables are reported as candidates for variables in the network of interest (cyan and blue curves). The *region of steep increase/decrease* is defined as the region where the piece-wise linear approximation is not constant (indicated by black line segments around the inflection points).

It is key to notice that in Fig. 1a, there is a change in the number of steady states. In phenotype 1 there is only one (stable) steady state: The state where the organism has its appendage (even after a cut, the appendage regenerates). In phenotype 2 there are two (stable) steady states: In addition to the steady state observed in phenotype 1, there is also the steady state where the organism does not have its appendage. This second state is stable because once the organism loses an appendage (after a cut), it will not regenerate. Thus, there is a bifurcation at *E* = *E*_0_ and we expect to see a “bifurcation-like” behavior of the response curves of the network variables involved in the change of phenotype. These variables (cyan and blue in the schematic) will presumably be the ones directly involved in the process of interest (in this case, in the process of regeneration). We remark that this approach does not need a model of the network. Since these changes in phenotype are observed at steady state, we assume that the response data is at steady state.

### 2.2 Toy model

To illustrate the approach in more detail, we use a toy model which was constructed with the following considerations.

1. The external stimulus (*E*) may not affect directly the variable of interest (say, *x*_2_), so we considered the presence of an intermediate variable (*x*_1_) between the external stimulus and the variable of interest.
2. The variable of interest may not only affect the phenotype (*y*), but also other variables in the network. So we considered the presence of a variable *x*_3_ downstream of the variable of interest.
3. The external input may affect other variables that are not (directly) connected to the variable of interested. So we considered the presence of a variable *x*_4_ affected by the external input such that it does not affect the variable of interest and is not affected by the variable of interest. Furthermore, *x*_4_ may have its own bifurcation-like behavior for some value of *E*.
4. There may be variables that are not affected by the external input neither directly nor indirectly. The response curves would be constant (with respect to *E*) in that case, so there was no need to consider such variables.

With these considerations we designed a toy network with 4 nodes, an external variable, and an output variable (Fig. 2a). The external variable, *E*, corresponds to external perturbations. The output variable, *y*, corresponds to a measure of the phenotype. The implementation of the network is as follows and will only be used to generate the data.

**Figure 2:**
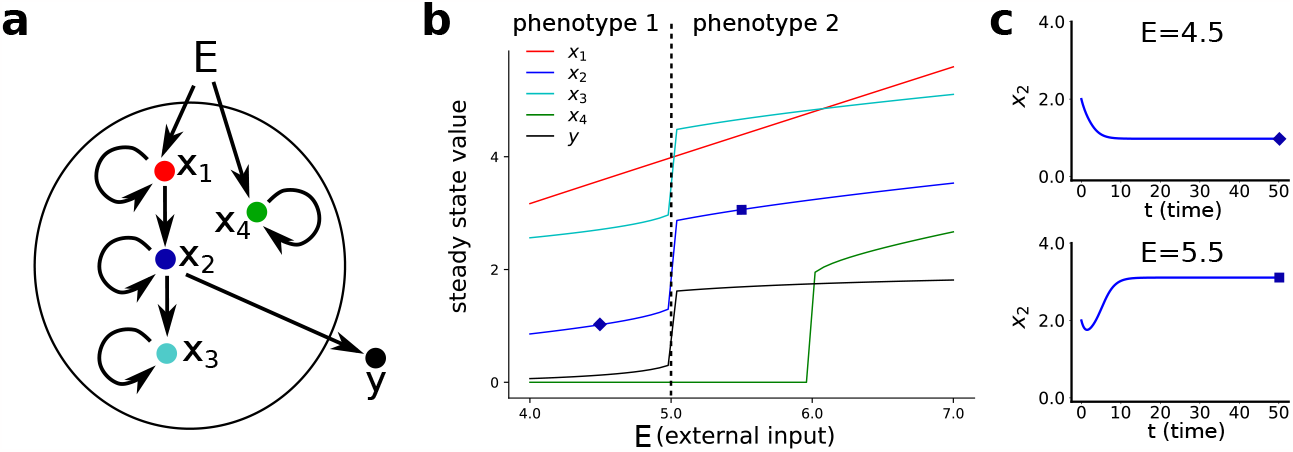
Small network. (**a**) Toy network subject to external perturbation. (**b**) Effect of external perturbation on network variables. The behavior of *x*_2_ and *x*_3_ are compatible with the transition between phenotypes at *E* = 5. The markers (diamond and square) indicate the values of the external input that correspond to the plots in panel (c). (**c**) Trajectory of *x*_2_ showing that a steady state has been reached. The value of *x*_2_ for this steady state is the value used in panel (b) at the corresponding marker. The initial condition for the simulations is (2,2,2,2), and the value of the parameters are *a*_1_ = 0.8, *a*_2_ = 2.4, *a*_3_ = 0.9, *a*_4_ = 0.41, *b*_2_ = 0.25, *b*_3_ = 2, *θ*_1_ = 1, *θ*_2_ = 2.1, *θ*_3_ = 2, *θ*_4_ = 1.4, *n* = 4, *a* = 2, *θ* = 2, *m* = 4

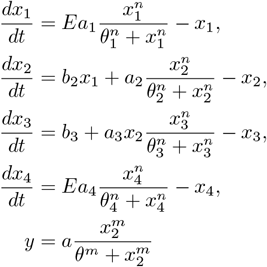

In this example, we refer to *phenotype 1* as the capability of the network to reach a steady state where the output variable is *y <* 1 (“low”) from *x* = (2, 2, 2, 2). If the network cannot achieve this, then we say it has phenotype 2. This particular setup is inspired by biological situations where a desired steady state exists (e.g. regeneration, apoptosis), but external inputs can disrupt the network in a way that makes that steady state unreachable (e.g. due to external chemicals, mutations).

For each value of the external stimulus, *E*, the data collected will be the steady state value reached from the initial condition (2, 2, 2, 2). We highlight that the only data available to use is the value of the variables in the network, the value of the external input, and the phenotype (whether *y <* 1). We can see a bifurcation-like behavior at different values of *E* (Fig. 2b), but since the transition between the phenotypes of interest happens at *E* = 5, we search for variables that exhibit a bifurcation-like behavior at that value of *E*, namely, *x*_2_ and *x*_3_. Variable *x*_2_ is indeed responsible for the change in phenotype (it directly affects *y*). However, *x*_3_ does not affect *y* and it was identified because it was downstream of *x*_2_. This is an important point that one has to take into consideration. The variables identified by this method are candidates, since downstream variables may also be identified. Importantly, note that for this approach to work, the external perturbation does not need to affect the subnetwork or module of interest directly. In the example, the perturbation goes through *x*_1_ before passing to *x*_2_ and our approach still works.

For response data that is continuous, one can compute the sensitivities with respect to *E* to see where there is a sudden change in the response. For data that is discrete and noisy, we can use the following algorithm.

### 2.3 Pseudocode

Here we present the pseudocode that we will use for discrete data.

#### Algorithm 1

Selecting variables of interest

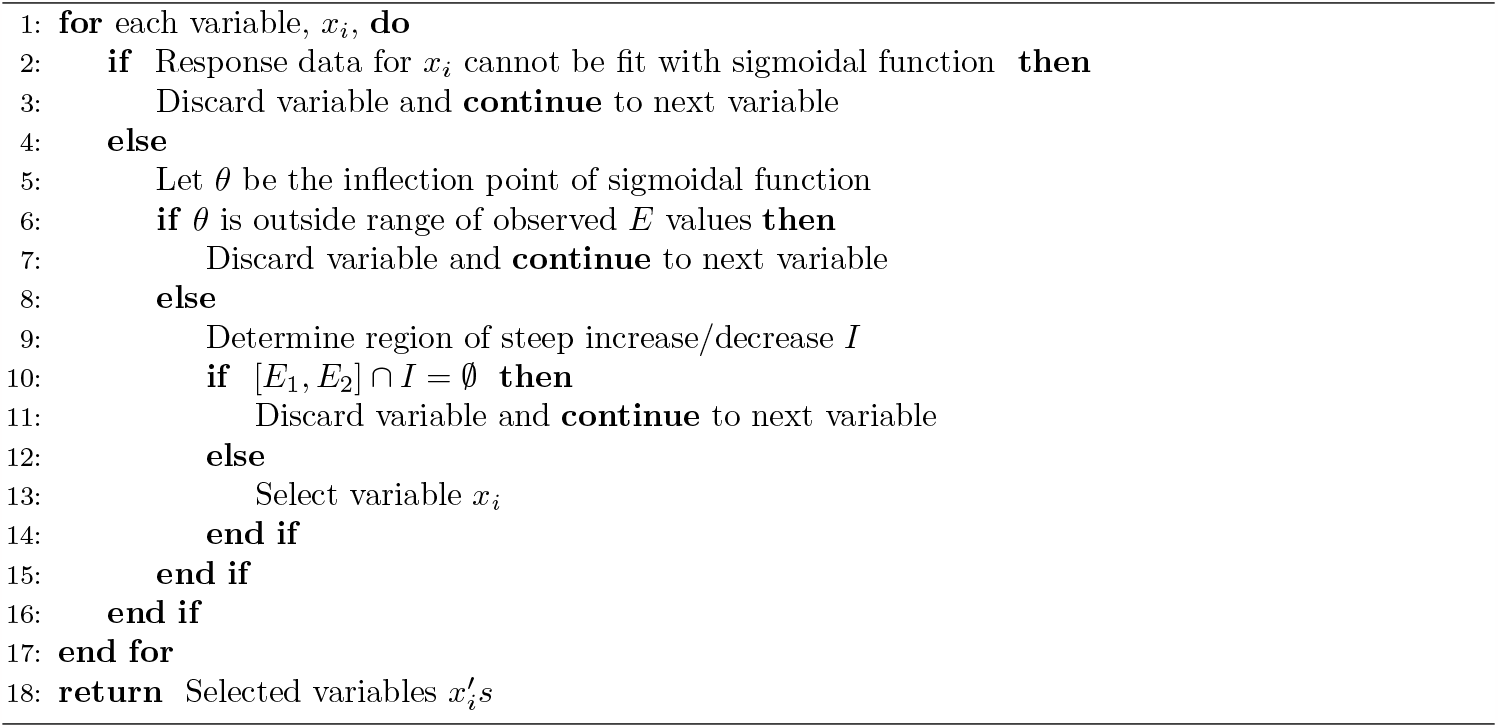

Steps 6-7 require further comment. If the data has a linear fit on the range of values of *E* observed in the data, [*E*_*low*_, *E*_*max*_], then it can be fit with a sigmoidal curve by artificially stretching it so that it looks linear on [*E*_*low*_, *E*_*max*_]. In these cases the inflection point will often be outside the range, and hence we discard those variables. If we need to discard variables that look linear or not sigmoidal enough, but still have their inflection point in [*E*_*low*_, *E*_*max*_], in Step 2 we can also impose a lower bound on the parameter that controls steepness of the fit.

We now show how to find the regions of steep increase/decrease for Step 9. First, we fit the discrete data with a sigmoidal function, *y* = *S*(*E*), Fig. 3a. Also, we find the tangent line to *S* at the inflection point *θ*, namely *y* = *S*^*′*^(*θ*)(*E* −*θ*)+*S*(*θ*), Fig. 3b. Then, we find the intersection of the tangent line and the horizontal line *y* = *a*, where *a* = lim_*E*→−∞_ *S*(*E*). Similarly, we find the intersection of the tangent line and the horizontal line *y* = *b*, where *b* = lim_*E*→−∞_ *S*(*E*). This results in the region 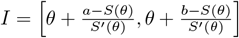, which we call the region of steep increase/decrease.

**Figure 3:**
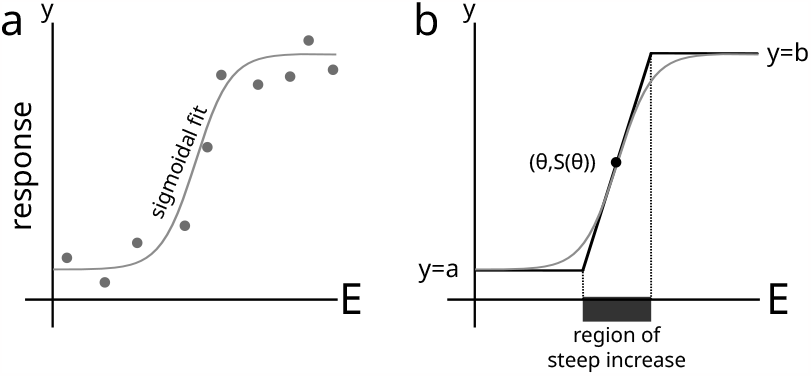
Computing the region of steep increase/decrease. (**a**) We first fit the data to a sigmoidal function, *y* = *S*(*E*) (if there is no good fit, the variable is discarded). (**b**) The region of steep increase/decrease is the interval where the piece-wise linear approximation is not constant.

For example, for the sigmoidal function 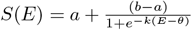 we have that 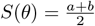 and 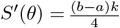. Then, the region of steep increase/decrease is 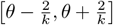.

## 3 In Silico Testing

### 3.1 Continuous data with no noise

We first test our approach on an *in silico* network with 24 variables such that it has an external input (*E*) and an output variable (*y*) which will determine the phenotype. Our goal is for our approach to return the variables of interest, with the caveat that downstream variables will also be returned.

As with the toy model, this network was designed with the following 4 considerations: (1) There is a subnetwork between the external stimulus and the subnetwork of interest, (2) the subnetwork of interest affects a downstream subnetwork, (3) there is a subnetwork affected by the external input that does not interact with the subnetwork of interest and has its own bifurcation-like behavior for some value of *E*, (4) any subnetwork that is not affected in any way by the external input would have a constant response curve, so it was not needed. With these considerations we constructed a network with 24 variables such that each of the 4 subnetworks mentioned above has 6 variables. Each subnetwork was designed as a continuous version of a qualitative model of the lac operon [9]. The connectivity of the network will be shown in Fig. 7b, but our approach starts with the assumption that the connectivity is unknown.

We used the synthetic network governed by the following differential equations.

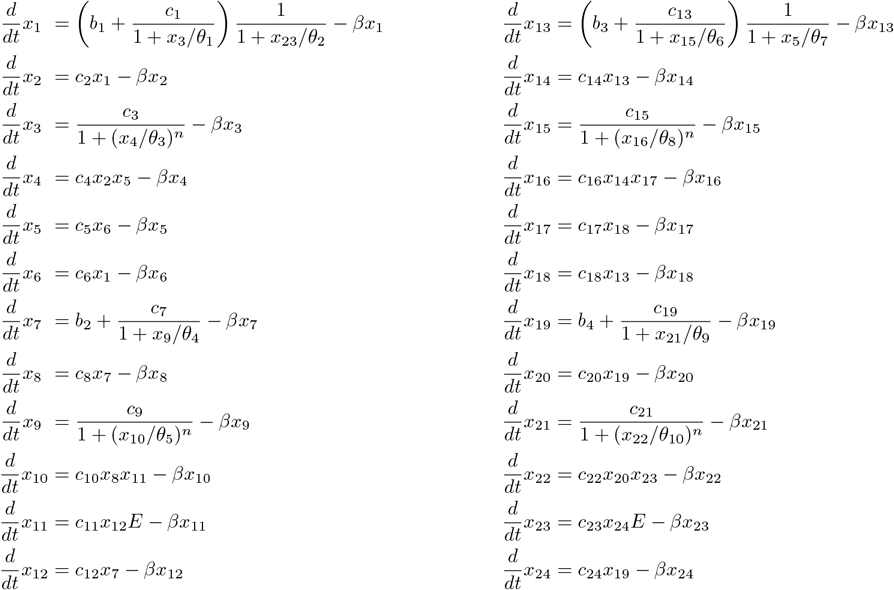

with output variable 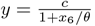.

In this example phenotype 1 refers to the capability of the system to reach a steady state where *y* is low, starting from the initial condition (1,1,0,1,1,1,1,1,0,1,1,1,1,1,0,1,1,1,1,1,0,1,1,1); and phenotype 2 is when the network cannot do this. We highlight that neither the wiring diagram nor the equations are used by the method, Fig. 4a. The method only needs response curves as its input and does not need prior information about the type of functions or the dependencies between variables. In this sense, the method is model-free. By increasing the external input, we find that the phenotype changes from 1 to 2 at *E* = 0.5, Fig. 4b. Thus, we are interested in variables whose response curves show a bifurcation-like behavior at *E* = 0.5.

**Figure 4:**
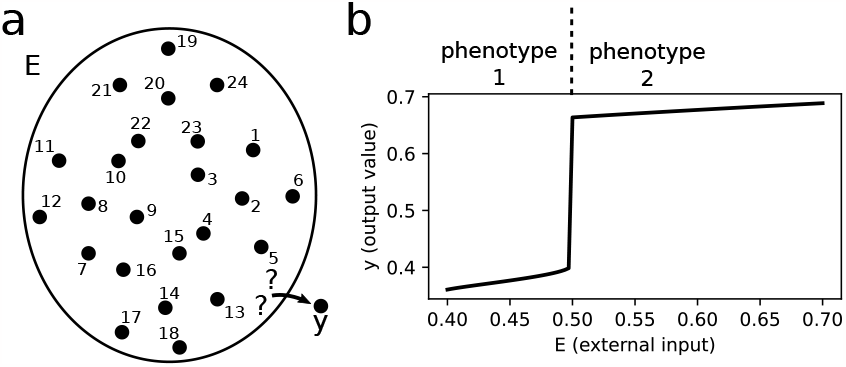
Network with change in phenotype. (a) Set of variables of in silico network subject to an external stimulus. Some variables directly affect the phenotype (encoded by variable *y*). The connectivity of the network is unknown. (b) Phenotype for different values of the external input. There is a change around *E* = 0.5 from phenotype 1 to phenotype 2.

Fig. 5 shows the response curves. We can see that some variables exhibit an abrupt change at *E* = 0.5. This can also be visualized by plotting the the sensitivities 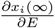 (we use *x*_*i*_(∞) to denote the value of *x*_*i*_ at steady state). Since the method would not have access to the model, we use a simple first-order finite difference. Later we will see how one can deal with data with limited data points (Subsection 3.2). Fig. 6 shows the sensitivities and variables of interest are those that have a peak or a valley at *E* = 0.5. In any case, the candidate variables are *x*_1_, *x*_2_, *x*_3_, *x*_4_, *x*_5_, *x*_6_, *x*_13_, *x*_14_, *x*_15_, *x*_16_, *x*_17_, *x*_18_.

**Figure 5:**
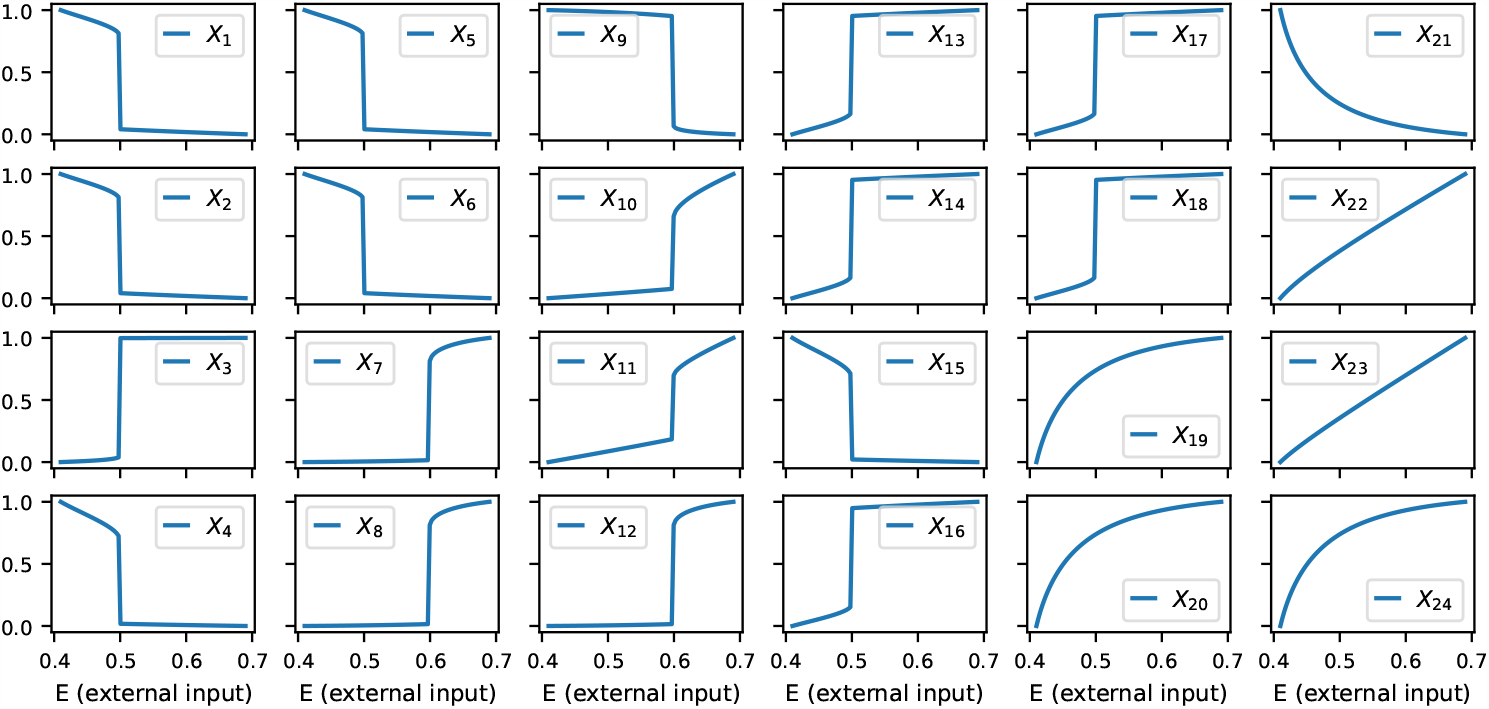
Response curves (normalized). Steady state value of the network variables for different values of the external input. Variables that exhibit an abrupt change around *E* = 0.5 are candidates to directly affect the phenotype of interest. The initial condition was (1,1,0,1,1,1,1,1,0,1,1,1,1,1,0,1,1,1,1,1,0,1,1,1), and the parameter values are *β* = 1, *b*_1_ = 0.011, *b*_2_ = 0.2, *b*_3_ = 0.3, *b*_4_ = 0.1, *c*_1_ = 0.7, *c*_2_ = 0.25, *c*_3_ = 1, *c*_4_ = 2, *c*_5_ = 4, *c*_6_ = 0.5, *c*_7_ = 0.7, *c*_8_ = 0.25, *c*_9_ = 1, *c*_10_ = 2, *c*_11_ = 2, *c*_12_ = 0.5, *c*_13_ = 0.525, *c*_14_ = 0.25, *c*_15_ = 100, *c*_16_ = 2, *c*_17_ = 4, *c*_18_ = 0.5, *c*_19_ = 0.7, *c*_20_ = 0.25, *c*_21_ = 1, *c*_22_ = 2, *c*_23_ = 4, *c*_24_ = 0.5, *θ*_1_ = 0.4, *θ*_2_ = 2, *θ*_3_ = 0.1, *θ*_4_ = 0.4, *θ*_5_ = 0.1, *θ*_6_ = 40, *θ*_7_ = 4, *θ*_8_ = 0.1, *θ*_9_ = 0.4, *θ*_10_ = 0.1, *n* = 4, *c* = 1, *θ* = 0.15.

**Figure 6:**
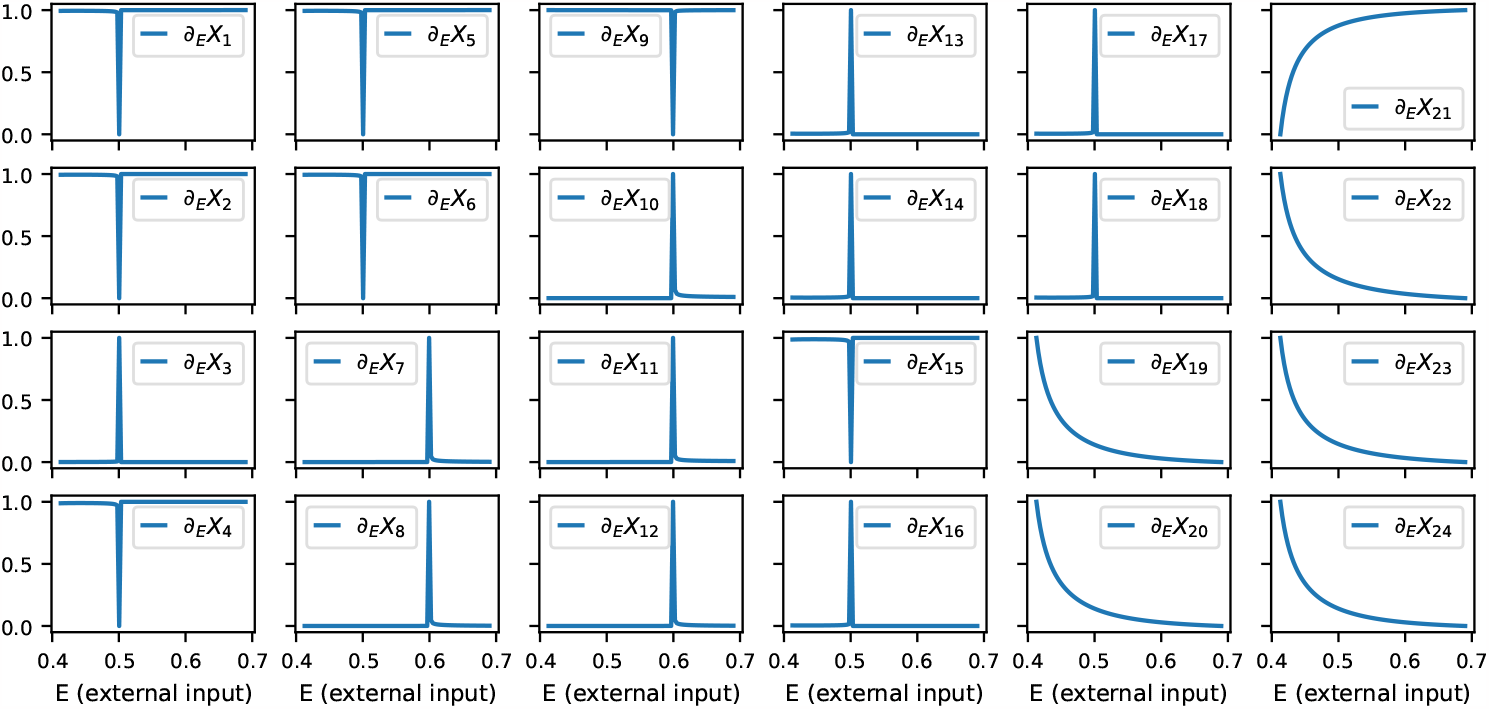
Sensitivity curves (normalized). By numerically computing 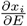 we can identify the variables to be reported by a peak around *E* = 0.5.

Fig. 7a shows all variables in the network with the reported variables highlighted. As validation, Fig. 7b shows the connectivity of the network. We can see that indeed the variables in the subnetwork or module that affects the phenotype directly was recovered. Also, consistent with the toy example in Section2, the downstream variables *x*_13_, *x*_14_, *x*_15_, *x*_16_, *x*_17_, *x*_18_ are also reported. Note that although the effect of the external input must pass through an upstream subnetwork or module (*x*_19_ through *x*_24_), the method still works. This is important, since in applications it is unknown *a priory* if an external input affects the variables of interest directly.

**Figure 7:**
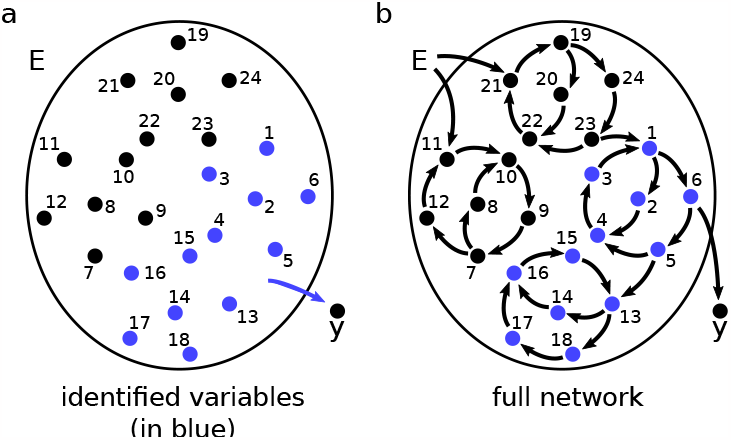
Variables reported by the proposed method. (a) Potential variables that may directly affect the phenotype are in blue. (b) The true network, for comparison. The method recovers all variables that affect directly the phenotype (encoded by *y*) and, as expected, it also reports all downstream variables.

### 3.2 Data with low resolution and noise

In practice, experiments may only be performed for a limited number of values of the external input. So, we now use response data that resembles real experimental data to test our approach given in Algorithm 1.

Suppose that data was generated using the network in Subsection 3.1 with the values of *E* = 0.40, 0.44, 0.48, 0.52, 0.56, 0.60, 0.64, 0.68, Fig. 8. Since the data is discrete, it is unknown where exactly the transition between phenotypes occurs; we only know it happens between 0.48 and 0.52. That is, [0.48, 0.52] is the transition region.

**Figure 8:**
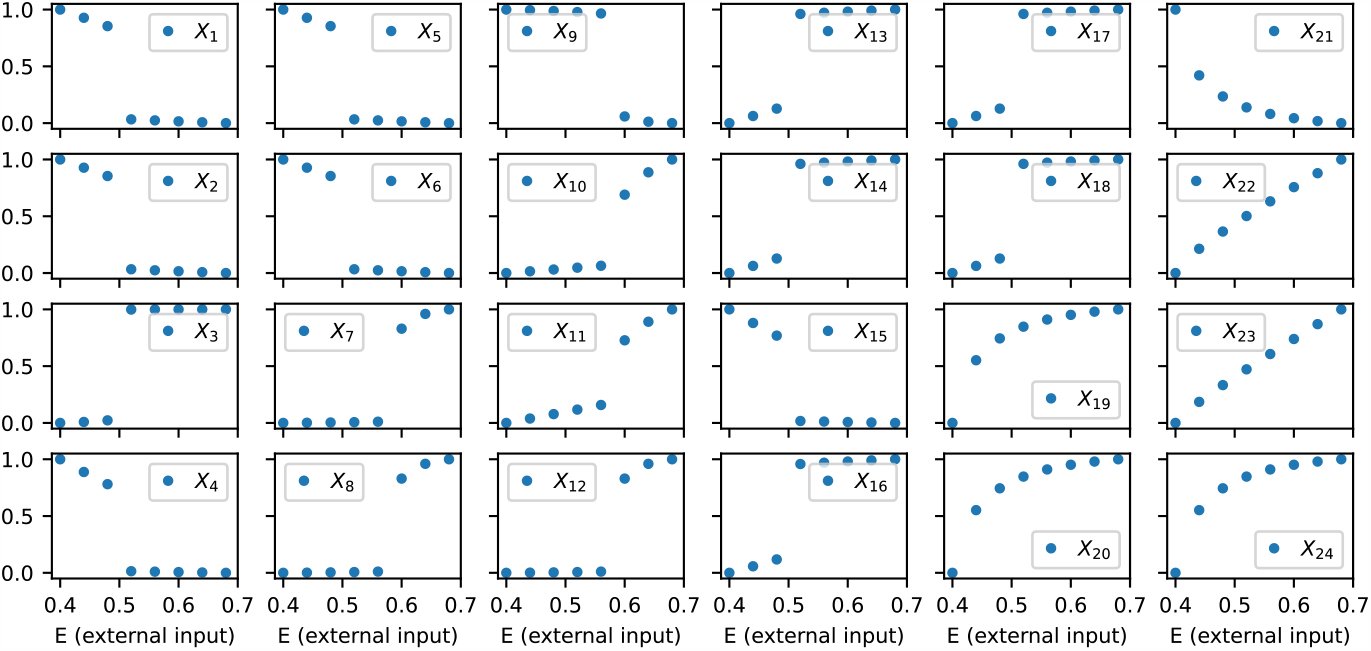
Low resolution response curves. Due to experimental constraints/costs or biological limitations data is not continuous.

First, we fit the response curves to a sigmoid function, 1. Here sigmoid means a class of functions that has an inflection point and that one of the parameters controls how steep the function is. We used the function

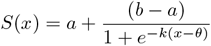

where *a* and *b* are the saturation values at low *x* and large *x*, respectively; *θ* is the inflection point and also the point where 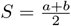; and *k* is the parameter that determines how steep the transition is around the inflection point.

Fig. 9 shows the results of using our approach. The only variables whose region of steep increase/decrease overlaps the transition region are *x*_1_, …, *x*_6_, *x*_13_, …, *x*_18_ (Fig. 7a). Other variables were discarded by our method due to not having a good sigmoidal fit or because there was no overlap between the regions.

**Figure 9:**
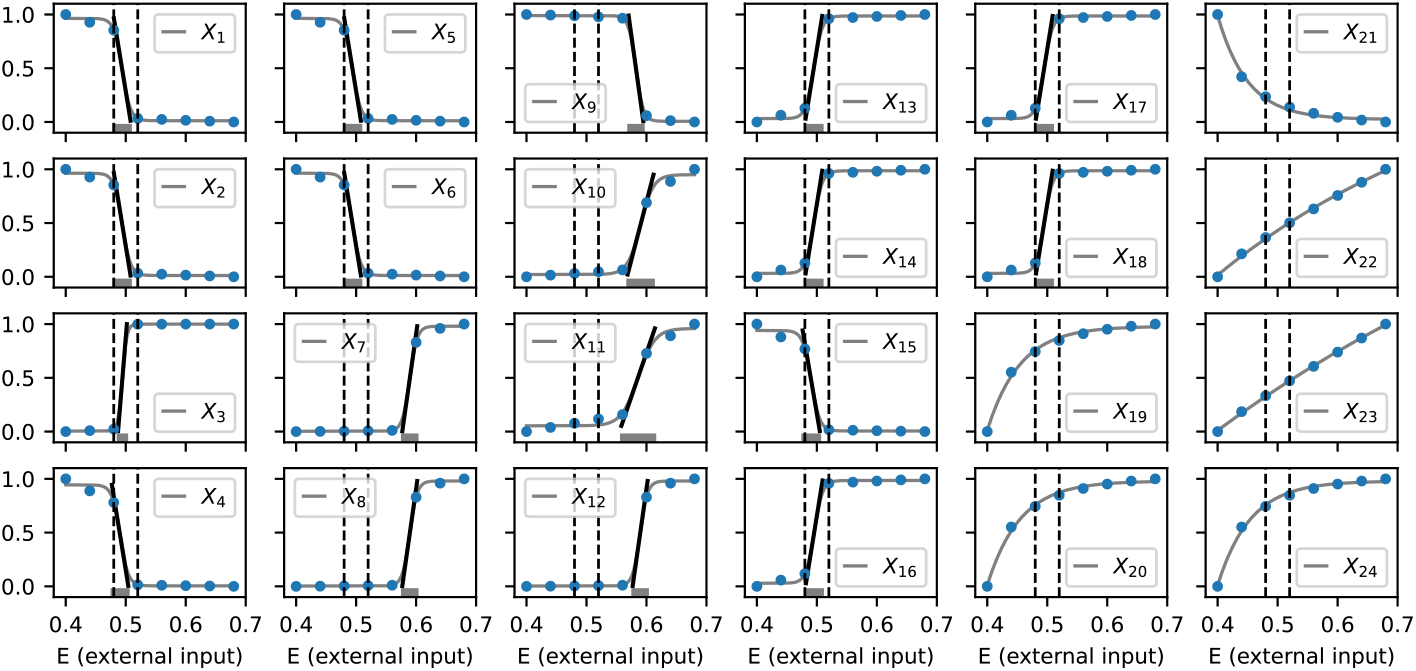
Selecting variables from discrete data. Our approach finds that variables *x*_1_, …, *x*_18_ can be fit with sigmoidal functions as required by Algorithm 1. For variables *x*_1_, …, *x*_6_, *x*_13_, …, *x*_18_ the transition region (between dashed lines) overlaps with the region of steep increase/decrease (indicated by gray rectangles). The output of the algorithm coincides with the case of continuous data (Fig. 7). Note: Sigmoidal fits can be obtained for variables *x*_19_, …, *x*_24_ (gray curves), but such variables are discarded since the inflection point is outside the window (Algorithm 1).

We also studied the effect of noise, Fig. 10. We added noise to each value of the response data using a normal distribution whose standard deviation was used as the noise level. We compared the noisy case with the noiseless case, so we considered *x*_1_, …, *x*_6_, *x*_13_, …, *x*_18_ as the variables to be reported (positives) and *x*_7_, …, *x*_12_, *x*_19_, …, *x*_24_ as variables to be discarded (negatives).

**Figure 10:**
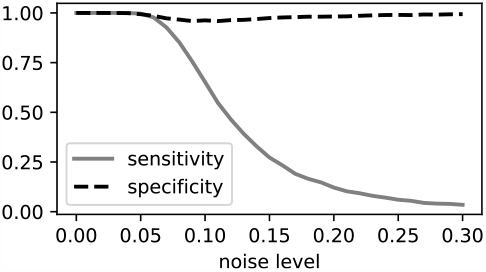
Sensitivity and specificity v.s. noise (the noiseless case is considered as the truth and the noise is given by a normal distribution with standard deviation given by the noise level). For small noise, both the sensitivity and the specificity are high. That is, all 12 variables *x*_1_, …, *x*_6_, *x*_13_, …, *x*_18_ are reported and the rest are excluded. Then, as noise increases, the number of variables reported decreases and eventually all of them are excluded.

For very low noise, both the specificity and the sensitivity were high. As noise increases, the sensitivity decreases until it eventually approaches zero. On the other hand, the specificity remains high for different noise levels. These findings can be explained as follows. As noise increases, less variables will have a sigmoidal fit, until eventually all variables are discarded. Also, variables that were discarded in the noiseless case have little chance to have a sigmoidal fit when noise is added and hence, they keep being discarded.

## 4 Application

We applied our variable selection method to a data set that was collected during salamanders (*ambystoma mexicanum*) tail regeneration [11]. The experimental setup consisted of two conditions: (1) the control case where under normal conditions the regeneration process was completed after amputating the tail, and (2) the intervention case where an HDAC class I chemical inhibitor (romidepsin) was applied after amputation to evaluate dose-dependent effects on regeneration and gene expression. For romidepsin concentrations 0, 0.01, and 0.05 *µ*M the regeneration process completed after tail amputation; we refer to this as phenotype 1. For romidepsin concentrations 0.5, 1.0, 5.0, and 10.0 *µ*M the regeneration process was inhibited after tail amputation; we refer to this as phenotype 2. Thus, increasing chemical concentration induced a change in phenotype above and below a transition region [0.05 *µ*M,0.5 *µ*M], Fig. 11. The response data in these experiments is gene expression measured for each concentration level of the chemical.

**Figure 11:**
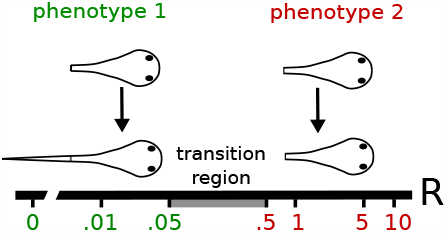
Experimental setup to measure phenotypic change in axolotls. After tail amputation, individuals were treated with different concentrations of the chemical romidepsin for 12 hours. With increasing concentration, we see a transition from phenotype 1 (regeneration is completed) to phenotype 2 (regeneration fails).

Romidepsin not only induced a change in regenerative outcome at 7 days post-amputation, it also affected gene expression at an earlier timepoint (12 hr post amputation). For our analysis, we selected 100 genes that were differentially expressed in the control and intervention cases [11]. The response data for these genes can be found in the supplementary material (excel file from Jupyter notebook).

We used Algorithm 1 to analyze these dose response data, first fitting genes (as response curves) with a sigmoidal fit, and then calculating an interval of steep increase/decrease using the inflection point and the tangent line to this point (Fig. 3). Then, for each gene, we checked if its corresponding interval intersected the transition region (i.e., [0.05 *µ*M,05 *µ*M] for this chemical).

We identified the following 12 genes: Axin1, Cbx4, Ccnl2, Cyp26b1, Dusp5, Eif4a3, Has2, Id1, Mas1, Nr4a1, Socs1, Sqstm1. The fits and errors (rescaled distance between data and fit) of these genes are provided in Figs. 12-13.

**Figure 12:**
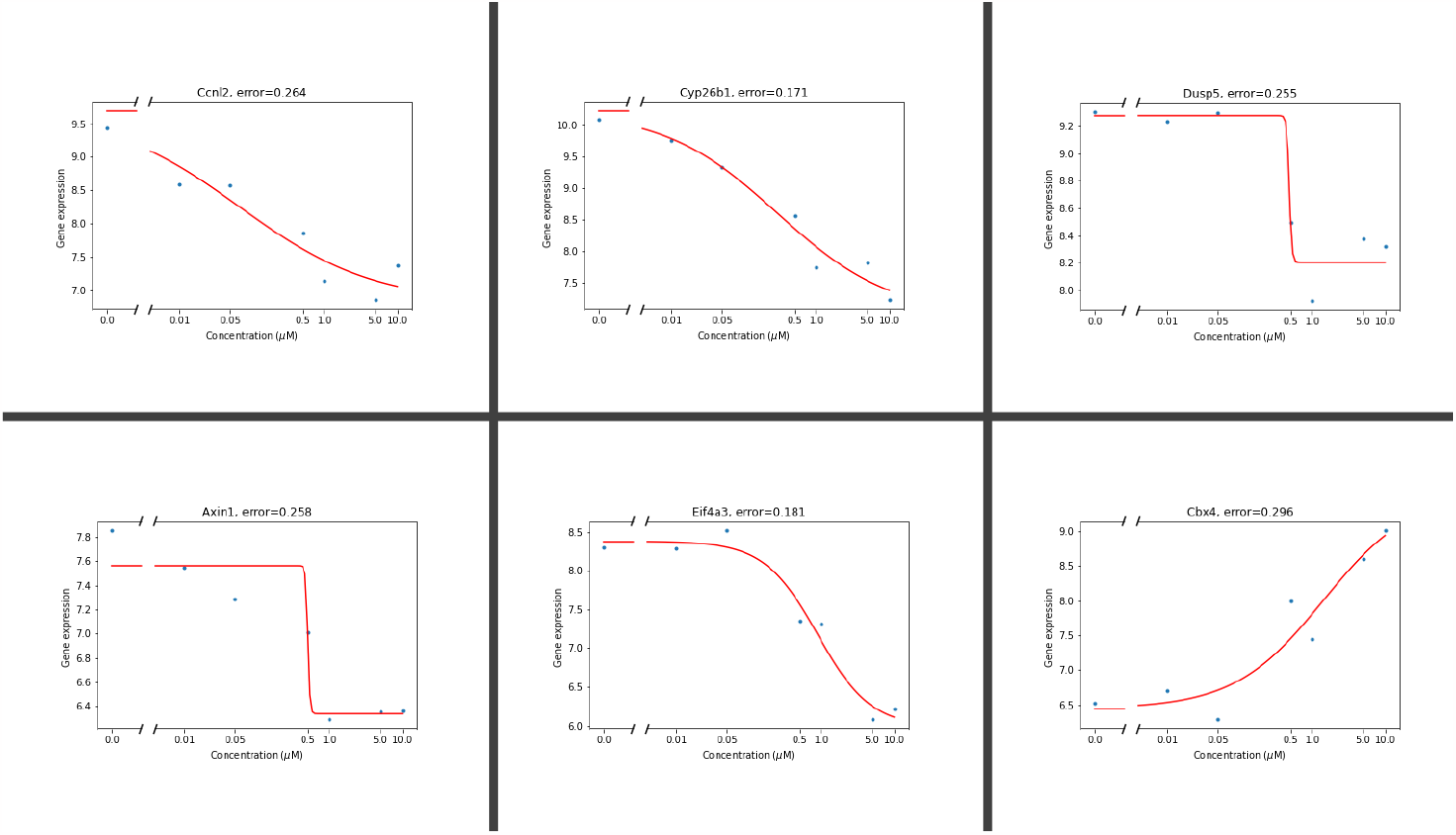
Selected variables for 12 HPA (hours post amputation).

It is hard to measure the accuracy of the method with real data because we don’t know which genes are directly affected by the chemical to alter tail regeneration. Thus, we may be missing some genes that might have been responsible for the block in regeneration. Nevertheless, Has2 (last gene in Fig. 13) was shown to be required for successful tail regeneration using CRISPR-Cas9 knockdown [11]. Some genes that were not selected included Cited2, Lep, Smad7 and GOs2. Consistent with the exclusion of these genes, F1 Cited2 knockout embryos complete regeneration after tail amputation (unpublished data).

**Figure 13:**
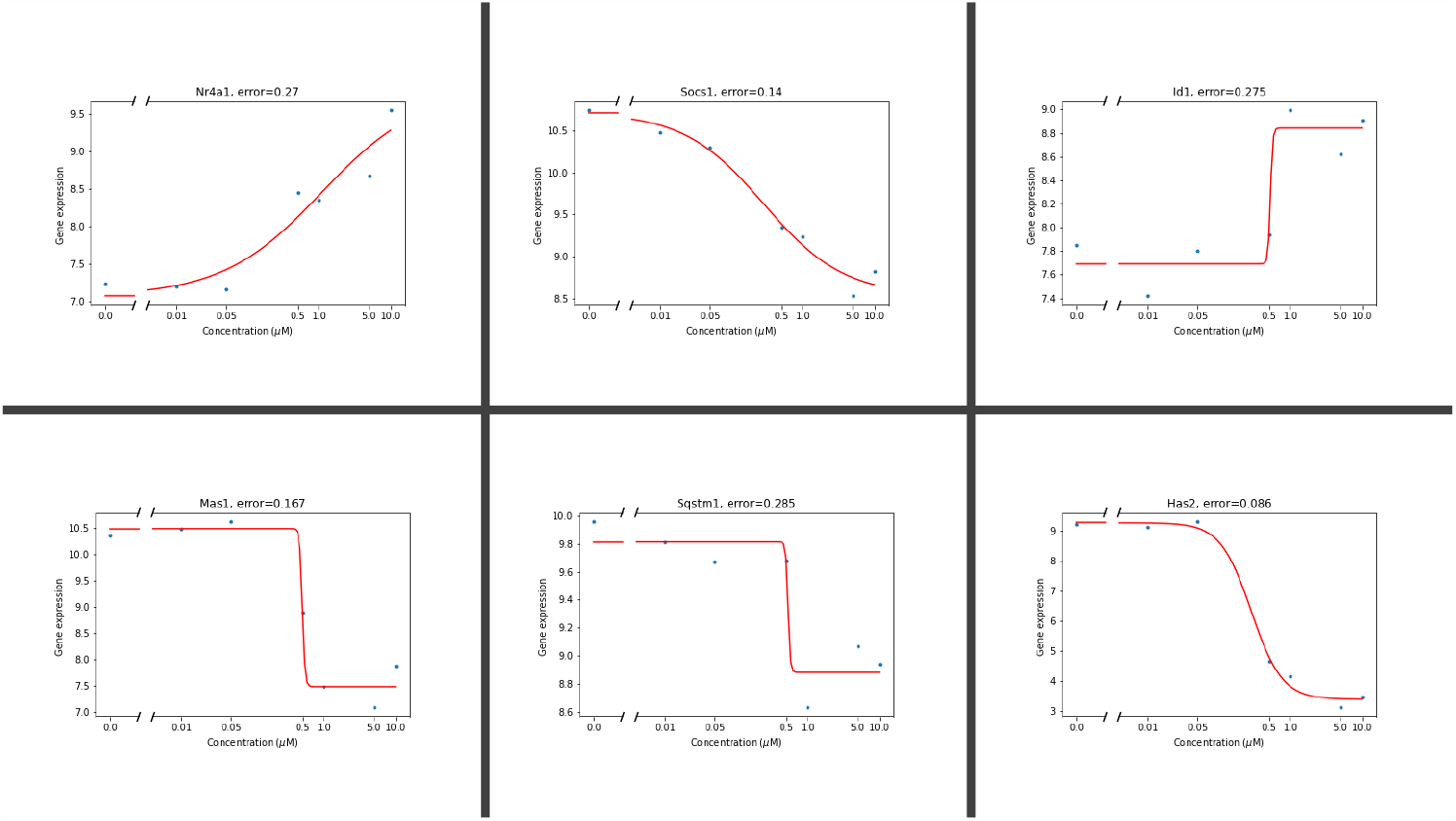
Selected variables for 12 HPA.

## 5 Discussion and Conclusion

In this paper we presented a method that uses dose response data to identify variables that are associated with complex phenotypic changes. Within the contexts of development and aging, an individual’s phenotype may change in response to a rising concentration of a chemical, such as a hormone that naturally triggers metamorphosis or puberty, or a chemical that transforms a normally functioning cell into a cancerous cell. Such chemicals can be manipulated within dose response experiments to identify critical thresholds that regulate phenotypic transitions. Our method efficiently identifies candidate variables from continuous response data corresponding to alternative phenotypic outcomes. For data that is discrete and noisy, our approach also identifies candidate variables by first fitting the data with sigmoidal functions to identify steep increases/decreases of the responses. Then, the variables that we identify are those that present abrupt transcriptional changes that overlap with the region where the phenotypic outcome changes. We applied our approach to gene expression during salamanders tail regeneration and 12 candidate genes were selected that can be used for experimental investigation. One of these genes, Has2, has been shown to be required for tail regeneration.

An advantage of the method is that it is straightforward to implement and there is no need for targeted perturbations such as knockout/over-expression experiments. We also showed that model predictions are robust with respect to a small amount of noise.

It is important to mention that care is needed in the interpretation of the reported variables/genes. With a toy model and an in-silico model we showed that the method does not distinguish between variables that are affecting the process of interest directly and downstream variables. Thus, the reported variables need to be interpreted as candidates for further investigation using experimental approaches. One potential way to overcome this limitation is to test multiple chemicals that presumably elicit the same phenotypic outcome.

Another limitation of this method is that it requires several data points to obtain a good fit. Response data with several data points can be expensive to collect. Another possible issue regarding the fitting step is that response data may have an abrupt change at more than one concentration. For such data, in the fitting step of the algorithm one can use multi-phasic functions that can model more than one region of steep increase [3]. Nevertheless, the variables reported would still be the variables that have one of their abrupt changes at the value of the chemical where a change in phenotype occurs.

The method presented here assumes that the response data are at steady state. This assumption is hard to verify in practice and gene networks are known to operate dynamically [4]. However, for some chronic disease states or irreversible phenotypic outcomes, such as when cells differentiate in mammals [2] or an organism transitions during ontogeny to an irreversible developmental stage [8], the variables of interest may accurately detail steady state conditions. Additionally, if we consider different time points in addition to different chemical concentrations, our method may be extended to detect relevant variables from continuous data using response surfaces (instead of response curves) and from discrete data by deriving a region of steep change (instead of interval of steep increase/decrease).

## Notes

https://github.com/alanavc/id-vars-from-resp-data

### Competing Interest Statement

The authors have declared no competing interest.

